# Quantifying the contribution of recessive coding variation to developmental disorders

**DOI:** 10.1101/201533

**Authors:** Hilary C. Martin, Wendy D. Jones, James Stephenson, Juliet Handsaker, Giuseppe Gallone, Jeremy F. McRae, Elena Prigmore, Patrick Short, Mari Niemi, Joanna Kaplanis, Elizabeth Radford, Nadia Akawi, Meena Balasubramanian, John Dean, Rachel Horton, Alice Hulbert, Diana S. Johnson, Katie Johnson, Dhavendra Kumar, Sally Ann Lynch, Sarju G. Mehta, Jenny Morton, Michael J. Parker, Miranda Splitt, Peter D. Turnpenny, Pradeep C. Vasudevan, Michael Wright, Caroline F. Wright, David R. FitzPatrick, Helen V. Firth, Matthew E. Hurles, Jeffrey C. Barrett, on behalf of the DDD Study

## Abstract

Large exome-sequencing datasets offer an unprecedented opportunity to understand the genetic architecture of rare diseases, informing clinical genetics counseling and optimal study designs for disease gene identification. We analyzed 7,448 exome-sequenced families from the Deciphering Developmental Disorders study, and, for the first time, estimated the causal contribution of recessive coding variation exome-wide. We found that the proportion of cases attributable to recessive coding variants is surprisingly low in patients of European ancestry, at only 3.6%, versus 50% of cases explained by *de novo* coding mutations. Surprisingly, we found that, even in European probands with affected siblings, recessive coding variants are only likely to explain ~12% of cases. In contrast, they account for 31% of probands with Pakistani ancestry due to elevated autozygosity. We tested every gene for an excess of damaging homozygous or compound heterozygous genotypes and found three genes that passed stringent Bonferroni correction: *EIF3F*, *KDM5B*, and *THOC6*. *EIF3F* is a novel disease gene, and *KDM5B* has previously been reported as a dominant disease gene. *KDM5B* appears to follow a complex mode of inheritance, in which heterozygous loss-of-function variants (LoFs) show incomplete penetrance and biallelic LoFs are fully penetrant. Our results suggest that a large proportion of undiagnosed developmental disorders remain to be explained by other factors, such as noncoding variants and polygenic risk.

---

Genetic studies of rare diseases traditionally followed a phenotype-driven search for a shared genetic diagnosis in multiple individuals with a clinically similar presentation. Large-scale sequencing studies of more phenotypically heterogenous patients have inverted this process, and have demonstrated the power of unbiased, genotype-first discovery of new disease genes ^1–3^. As this approach to gene discovery becomes routine, it is also possible to use the same datasets to characterise the overall genetic architecture of such disorders. For example, we previously showed how analyses of one such cohort, the Deciphering Developmental Disorders study (DDD) ^4^, can discover new dominant disease genes ^1^, estimate the fraction of patients with a causal *de novo* mutation in both known and as-yet undiscovered dominant genes (40-45%), and make predictions about the population prevalence of such disorders ^5^.

Extending this characterisation to other modes of inheritance will help design future gene discovery studies and inform counselling about recurrence risk. It has been posited, based on extrapolations from the X chromosome, that there are potentially thousands of recessive genes yet to be discovered ^6^, which could imply that recessive genes explain a large fraction of undiagnosed rare disease cases. However, attempts to estimate the prevalence of recessive disorders in the population have been restricted to known disorders ^7^ or known pathogenic alleles ^8^. While families with multiple affected siblings or parental consanguinity are thought more likely to have a recessive disorder ^9^, there has been no systematic attempt to quantify the recessive burden using large-scale sequencing data and a robust statistical and computational genetic framework.

Here we describe an analysis of autosomal recessive coding variants in 7,448 exome-sequenced families from the British Isles, recruited as part of the DDD study. We previously developed a probabilistic method for identifying robust new recessive genes ^10^ significantly enriched for biallelic (homozygous or compound heterozygous) genotypes, an approach that stands in contrast to the heuristic filtering methods commonly applied ^11–17^. We extend this to estimate the overall burden of recessive causes in this cohort, and compare this between different groups of patients with British European or British Pakistani ancestry. Our approach overcomes drawbacks of previously published methods ^3,10^ that do not provide well-calibrated estimates of this overall exome-wide burden of recessive disease.

## Results

### Genome-wide recessive burden

We hypothesized there should be a burden of biallelic genotypes predicted either to cause loss-of-function (LoF) or likely damage to a protein. For each of three possible genotype configurations (LoF on both alleles, damaging missense on both alleles, or one of each), we compared the number of observed rare (minor allele frequency, MAF, <1%) biallelic genotypes in our cohort to the number expected by chance given the population frequency of such variants and the gene-specific fraction of individuals who are autozygous (see Methods). Because the expected number is sensitive to inaccuracy in population frequency estimates of very rare variants in broadly-defined ancestry groups like “Europeans” or “South Asians”, we focused our analysis on the largest two subsets of the cohort that had homogenous ancestry, corresponding in a principal components analysis using 1000 Genomes Project populations to Great British individuals and Punjabis from Lahore, Pakistan (Supplementary Figure 1). We refer to these subsets as being of European Ancestry or Pakistani Ancestry from the British Isles (EABI, PABI).

We evaluated three possible methods for calculating the expected number of biallelic genotypes, using synonymous variants as a control since we do not expect these to be involved in disease. Firstly, we used the non-Finnish Europeans and South Asians from the Exome Aggregation Consortium (ExAC) ^18^ to estimate the allele frequencies for EABI and PABI respectively, as we described previously ^10^. However, we found that the total observed number of biallelic synonymous genotypes was dramatically lower than the expected number (Supplementary Figure 2). This is due to a combination of differences in sequence coverage, quality control, and ancestry between DDD and ExAC, and the lack of phased, individual-specific data in ExAC needed to avoid double-counting variants on the same haplotype within a gene. Secondly, we considered a recently published approach ^3^ based on per-gene mutation rates. While it was reported to be well-calibrated for individual genes, this method produced a significant underestimate of the total expected number of synonymous biallelic genotypes (Supplementary Figure 2). Finally, we decided to use the phased haplotypes from unaffected DDD parents to estimate the expected number of biallelic genotypes henceforth (see Methods). With this method, the number of observed biallelic synonymous variants closely matched what we would expect by chance (ratio=0.997 for EABI and 1.003 for PABI; p=0.6 and 0.4) (Figure 1A). We observed no significant burden of biallelic genotypes of any consequence class in 1,366 EABI probands with a likely diagnostic *de novo* mutation, inherited dominant variant or X-linked variant, consistent with those probands’ phenotypes being fully explained by the variants already discovered. We therefore evaluated the recessive coding burden in 4,318 EABI and 333 PABI probands whom we deemed more likely to have a recessive cause of their disorder because they did not have a likely diagnostic variant in a known dominant or X-linked developmental disorder (DD) gene ^4^, or had at least one affected sibling, or >2% autozygosity. As expected due to their higher autozygosity (Supplementary Figure 3), PABI individuals had substantially more rare biallelic genotypes than EABI individuals (Figure 1). Ninety-two percent of the likely damaging rare biallelic genotypes observed in PABI samples were homozygous (rather than compound heterozygous), versus only 28% for the EABI samples. We observed a significant enrichment of biallelic LoF genotypes above chance expectation in both the EABI and PABI groups (~1.4-fold enrichment in each; p=3.5×10^−5^ for EABI, p=9.7×10^−7^ for PABI). We also observed a smaller enrichment of biallelic damaging missense genotypes which was nominally significant in the EABI group (p=0.03), as well as a significant enrichment of compound heterozygous LoF/damaging missense genotypes in the EABI group (1.4-fold enrichment; p=6×10^−7^). In the EABI group, the enrichments became stronger and more significant at lower MAF, but the absolute number of excess variants fell slightly (Supplementary Figure 4). Thus, plausibly pathogenic variants are concentrated at rarer MAF, but some do rise to higher frequencies.

**Figure 1:**
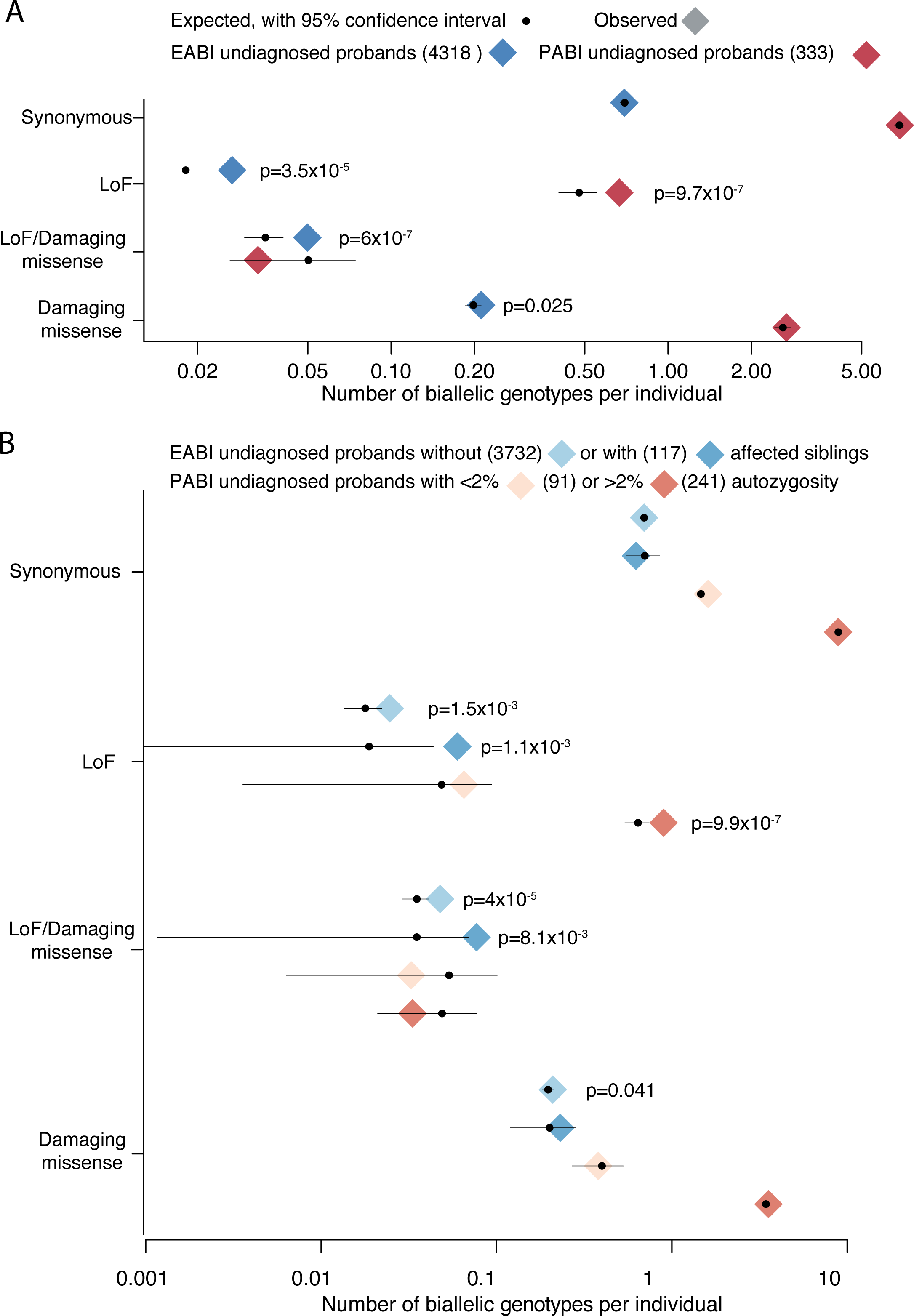
Number of observed and expected biallelic genotypes per individual for all genes in A) undiagnosed EABI and PABI probands, and B) different subsets of undiagnosed probands. Nominally significant p-values from a test of enrichment (assuming a Poisson distribution) are shown. The samples sizes are indicated in parentheses in the keys. Note that the set of probands with affected siblings includes only those whose siblings were also in DDD and appeared to share the same phenotype (see Methods).

We found that particular gene sets showed a higher burden of damaging biallelic genotypes. A set of 903 clinically-curated DD-associated recessive genes showed very strong enrichment of damaging biallelic genotypes (1.7-fold; p=6×10^−18^ for EABI and PABI combined). Indeed, 48% of the observed excess of damaging biallelic genotypes could be accounted for by these known genes. We also found a highly significant enrichment of damaging biallelic genotypes in 3371 genes annotated as having high probability of being intolerant of LoFs in the recessive state (pRec>0.9) ^18^ (1.2-fold; p=2×10^−8^), even after excluding the known recessive genes used to train the model (1.1-fold, p=4×10^−4^), and in 189 genes that were sub-viable when knocked out homozygously in mice ^19^ (1.8-fold; p=2×10^−3^). By contrast, we did not observe any recessive burden in 243 DD-associated genes that act by a dominant LoF mechanism, nor in genes predicted to be intolerant of heterozygous LoFs (probability of LoF intolerance, pLI, >0.9) in ExAC.

We developed a new method to estimate the proportion of probands who have a diagnostic variant in a particular genotype class (Methods). In contrast to our previously published approach ^5^, our new method accounts for the fact that a fraction of the variants expected by chance are actually causal; thus, it gives slightly higher estimates than we previously reported for the proportion of the cohort with causative *de novo* mutations. We estimated that 3.6% of EABI probands have a recessive coding diagnosis, compared to 49.9% with a *de novo* coding diagnosis. In the PABI subset, recessive coding genotypes likely explain 30.9% of individuals, compared to 29.8% for *de novo* coding mutations. The contribution from recessive variants was nearly four times as high in EABI probands with affected siblings than those without affected siblings (12.0% versus 3.2%), and highest in PABI probands with high autozygosity (47.1%) (Figure 2). In contrast, it did not differ significantly between PABI probands with low autozygosity and EABI probands. Supplementary Table 1 shows the 95% confidence intervals of these diagnostic fractions for different consequence classes in different sample subgroups. These estimates rely on another parameter, the proportion of genotypes in a particular class that are pathogenic (Supplementary Figure 5), but in fact, they are not very sensitive to this (see Methods).

**Figure 2:**
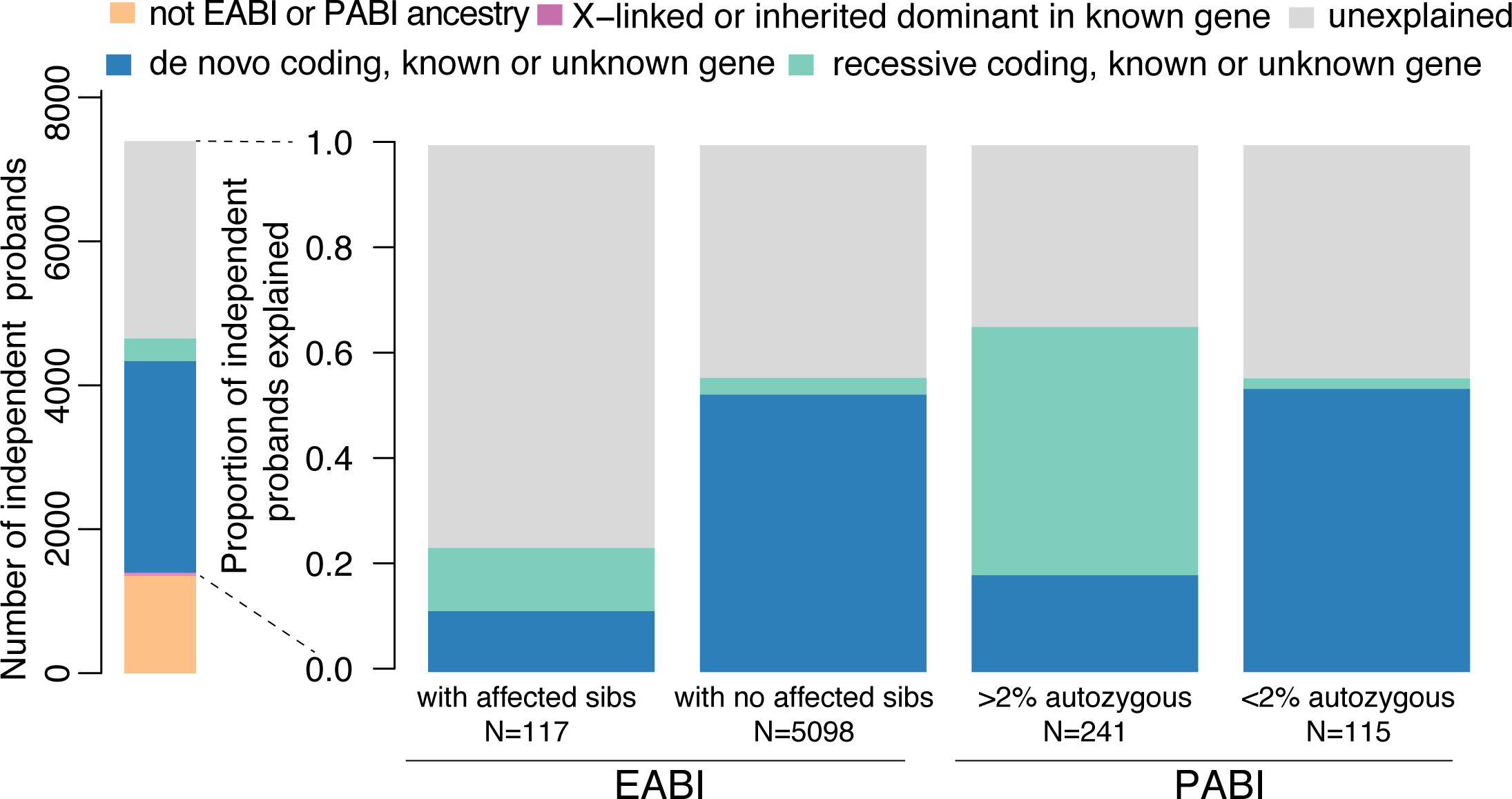
Left: number of independent trio probands grouped by diagnostic category. The inherited dominant and X-linked diagnoses include only those in known genes, whereas the proportion of probands with *de novo* and recessive coding diagnoses was inferred as described in the Methods. Right: the proportion of probands in various EABI and PABI subsets inferred to have diagnostic *de novo* coding mutations or recessive coding variants in known or as-yet-undiscovered genes.

### Discovery of new recessive disease genes

In order to discover new recessive genes, we next tested each gene in either EABI alone or EABI+PABI for an excess of biallelic genotypes. We tested four combinations of the consequence categories described above (Methods) because, in some genes, biallelic LoFs might be embryonic lethal and LoF/damaging missense compound heterozygotes might cause DD, but in other genes, including rare damaging missense variants in the analysis might drown out signal from truly pathogenic LoFs.

Three genes passed stringent Bonferroni correction (p<3.4×10^−7^, accounting for 8 tests for each of 18,630 genes), *EIF3F*, *KDM5B* and *THOC6*, of which the latter is an established recessive DD-associated gene ^20–22^. Thirteen additional genes had p-value<10^−4^ (Table 1), eleven of which are known recessive DD-associated genes, and the distribution of p-values for all known recessive DD-associated genes was shifted significantly lower than that of all other genes (Kolmogorov-Smirnov test; p<1×10^−15^; Supplementary Figure 6). Summary statistics for all genes are given in Supplementary Table 2. For six of the genes in Table 1, one or more families had affected siblings who shared the biallelic genotypes, supporting their pathogenicity. Patients with biallelic damaging genotypes in *THOC6*, *CNTNAP1*, *KIAA0586*, and *MMP21* were significantly more phenotypically similar to each other than expected by chance (phenotypic p-value given in Table 1). Taken together, these observations validate our gene discovery approach, and suggest that our genome-wide significance threshold is likely conservative.

**Table 1:** Genes enriched for damaging biallelic coding genotypes with p<1×10^−4^. The number of observed biallelic genotypes of different consequence classes is shown for the EABI and PABI probands. The lowest p-value out of the eight tests conducted is indicated, along with the details of the corresponding test and the p-value for phenotypic similarity for the relevant probands. For all genes except *VPS13B*, the lowest p-value was achieved using EABI alone. Known recessive DD genes from the DDG2P list are indicated (<u>http://www.ebi.ac.uk/gene2phenotype/)</u>.

| gene | Biallelic genotypes counts<br>for EABI (PABI if >0) |  |  | p-value -<br>genotype | p-value -<br>phenotype | Consequence class<br>for most significant<br>test | Note |
| --- | --- | --- | --- | --- | --- | --- | --- |
|  | LoF | LoF/<br>damaging<br>missense | damaging<br>missense |  |  |  |  |
| *EIF3F* | 0 | 0 | 5 | 1.2E-10 | 0.72 | damaging missense | two probands had another affected sibling, both of whom were homozygous for the same variant |
| *THOC6* | 0 | 1 | 3 | 4.4E-09 | 6.0E-05 | LoF + LoF/damaging missense + damaging missense | known recessive gene |
| *KDM5B* | 3 | 0 | 0 | 1.1E-07 | 0.53 | LoF | previously reported as dominant gene; a fourth proband is compound heterozygous for a splice variant and CNV |
| *CNTNAP1* | 2 (1) | 1 | 0 (1) | 1.8E-06 | 0.02 | LoF+LoF/damaging missense | known recessive gene |
| *KIAA0586* | 5 | 1 | 1 | 1.9E-06 | 0.05 | LoF | known recessive gene; two probands have affected sibs, and both share the variants |
| *NALCN* | 1 | 2 | 0 | 2.4E-06 | 0.37 | LoF+LoF/damaging missense | known recessive gene; one proband has an affected sib who shares the variants |
| *PIGN* | 0 | 3 | 1 | 2.5E-06 | 0.10 | LoF + LoF/damaging missense + damaging missense | known recessive gene; one proband has an affected sib who shares the variants |
| *ST3GAL5* | 0 (1) | 2 | 0 | 2.7E-06 | 0.09 | LoF+LoF/damaging missense | known recessive gene |
| *ATAD2B* | 1 | 1 | 0 | 3.6E-06 | 0.88 | LoF+LoF/damaging missense | one of our probands has an affected sib who shares the variants |
| *LZTR1* | 0 | 3 | 0 | 5.6E-06 | 0.06 | LoF+LoF/damaging missense | one of our probands has an affected sib who does not share both variants, so causality is dubious ;dominant missense mutations cause Noonan syndrome<sup>77</sup> |
| *LINS* | 2 | 0 | 0 | 8.2E-06 | 0.74 | LoF | one proband has affected sib who shares the variant; putative recessive gene<sup>78,79</sup> |
| *POLR1C* | 0 | 1 | 2 | 1.4E-05 | 0.42 | LoF + LoF/damaging missense + damaging missense | known recessive gene |
| *MMP21* | 0 | 2 | 0 | 1.4E-05 | 2.0E-03 | LoF+LoF/damaging missense | known recessive gene |
| *MAN1B1* | 1 | 1 | 0 | 1.5E-05 | 0.62 | LoF+LoF/damaging missense | known recessive gene |
| *VPS13B* | 2 (1) | 1 | 2 | 2.8E-05 | 0.05 | LoF | known recessive gene |
| *UBA5* | 0 | 2 | 1 | 3.7E-05 | 0.84 | LoF+LoF/damaging missense | known recessive gene |

We observed five probands with an identical homozygous missense variant in *EIF3F* (p=1.2×10^−10^) (ENSP00000310040.4:p.Phe232Val), which is predicted to be deleterious by SIFT, polyPhen and CADD. There were an additional four individuals in the DDD cohort who were also homozygous for this variant but who had been excluded from our discovery analysis: two were siblings of distinct index probands, one had a potentially diagnostic inherited X-linked variant in *HUWE1* (subsequently deemed to be benign since it did not segregate with disease in his family), and one had no parental genetic data available. All probands had European ancestry and low overall autozygosity, and none of them (apart from the pairs of siblings) were related (kinship<0.02). In the gnomAD resource of population variation (<u>http://gnomad.broadinstitute.org/</u>), this variant (rs141976414) has a frequency of 0.12% in non-Finnish Europeans, and no homozygotes were observed.

All nine individuals homozygous for the *EIF3F* variant had ID and six had seizures (Supplementary Table 3). Affected individuals for whom photos were available did not have a distinctive facial appearance (Supplementary Figure 7). Features observed in three or more unrelated individuals were behavioural difficulties and sensorineural hearing loss. One of these individuals was previously published in a case report ^23^. One patient had skeletal muscle atrophy (Supplementary Figure 7), which is only reported in one other proband in the DDD study. This is notable because in mice, *Eif3f* has been shown to play a role in regulating skeletal muscle size via interaction with the mTOR pathway ^24^. None of the other individuals were either assessed to have or previously recorded to have muscle atrophy.

*EIF3F* encodes the F subunit of the mammalian eIF3 (eukaryotic initiation factor) complex, a negative regulator of translation. The genes encoding eIF2B subunits have been implicated in severe autosomal recessive neurodegenerative disorders ^25^. The secondary structure, domain architecture and 3D fold of EIF3F is well conserved between species but sequence similarity is low (29% between yeast and humans) (Figure 3A). The highly conserved Phe232 side chain mutated in our patients is buried (solvent accessibility 0.7%) and likely plays a stabilising role, perhaps in conjunction with two other conserved aromatic amino acids (Figure 3B). The loss of the aromatic side chain in the Phe232Val variant would likely disrupt protein stability. Further work will be needed to understand how the Phe232Val variant affects EIF3F function, and how this causes DD.

**Figure 3:**
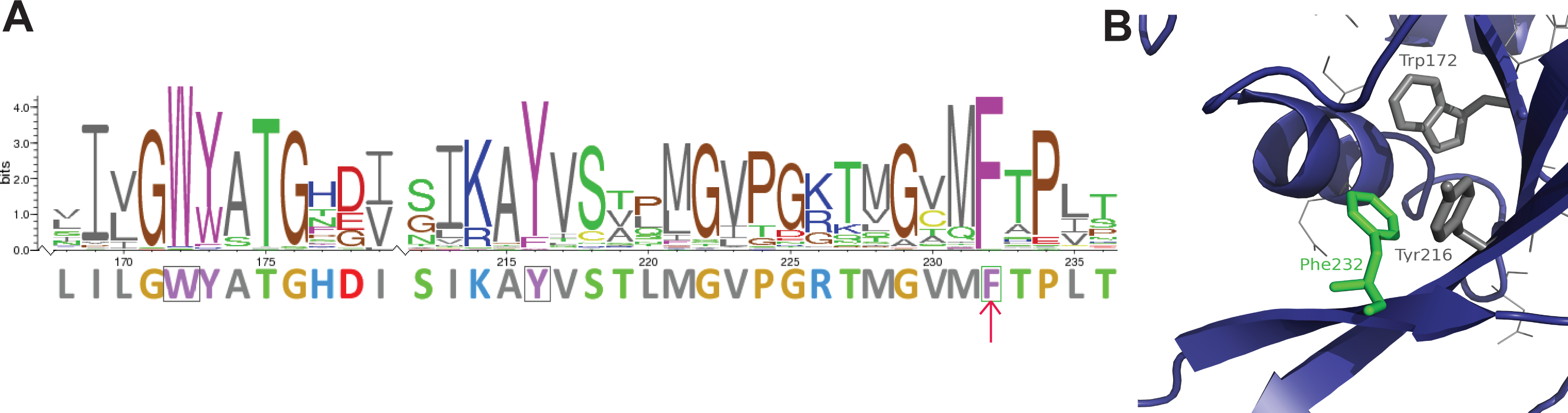
a) Section of the amino acid sequence logo for EIF3F where the strength of conservation across species is indicated by the size of the letters. The sequence below in fixed height characters represents the human EIF3F. Boxed characters are those aromatic residues conserved between humans and yeast and proximal in space to Phe232. b) Structure of the section of EIF3F containing the Phe232Val variant, highlighted in green. The blue backbone is from an X-ray structure of yeast 26S proteasome regulatory subunit RPN8 (PDB entry 4OCN), which is structurally virtually identical to human EIF3F (RMSD from PDB entry 3J8C:f <1Å). Amino acids conserved between yeast and human sequences as highlighted in panel a are shown in grey.

The second new recessive gene we identified was *KDM5B* (p=1.1×10^−7^) (Figure 4). *KDM5B* encodes a histone H3K4 demethylase. Other H3K4 methylases (*KMT2A, KMT2C, KMT2D, SETD1A*), demethylases (*KDM1A, KDM5A, KDM5C*), and two related reader proteins (*PHF21A, PHF8*) are known to cause neurodevelopmental disorders ^26–28^. Three probands had biallelic LoFs passing our filters, and we subsequently identified a fourth who was compound heterozygous for a splice site variant and large gene-disrupting deletion. Curiously, *KDM5B* is also enriched for *de novo* mutations in the DDD cohort ^5^ (p=5.1×10^−7^). Additionally, we saw nominally significant over-transmission of LoF variants from the parents, who were almost all unaffected (p=0.002 including all families; p=0.02 when biallelic trios were excluded; transmission-disequilibrium test). This suggests that heterozygous LoFs in *KDM5B* confer an increased risk of DD but are not fully penetrant, which is consistent with the observation of 22 LoF variants in ExAC (pLI = 0), very unusual for dominant DD genes. We considered the possibility that all the *KDM5B* LoFs observed in probands might be, in fact, acting recessively and that the probands with apparently monoallelic LoFs had a second coding or regulatory hit on the other allele. However, we found no evidence supporting this hypothesis (see Methods and Supplementary Figure 8), nor of potentially modifying coding variants in likely interactor genes. There was also no evidence from the annotations in Ensembl or GTex data (<u>https://gtexportal.org/home/</u>) that the pattern could be explained by some LoFs being evaded by alternative splicing or avoiding nonsense-mediated decay (Figure 4B). We ran methylation arrays to search for an epimutation that might be acting as a modifier in the apparently monoallelic LoF carriers, but found none (Supplementary Figure 9). Together, these different lines of evidence suggest that heterozygous LoFs in *KDM5B* are pathogenic with incomplete penetrance, while homozygous LoFs are, as far as we can tell, fully penetrant.

**Figure 4:**
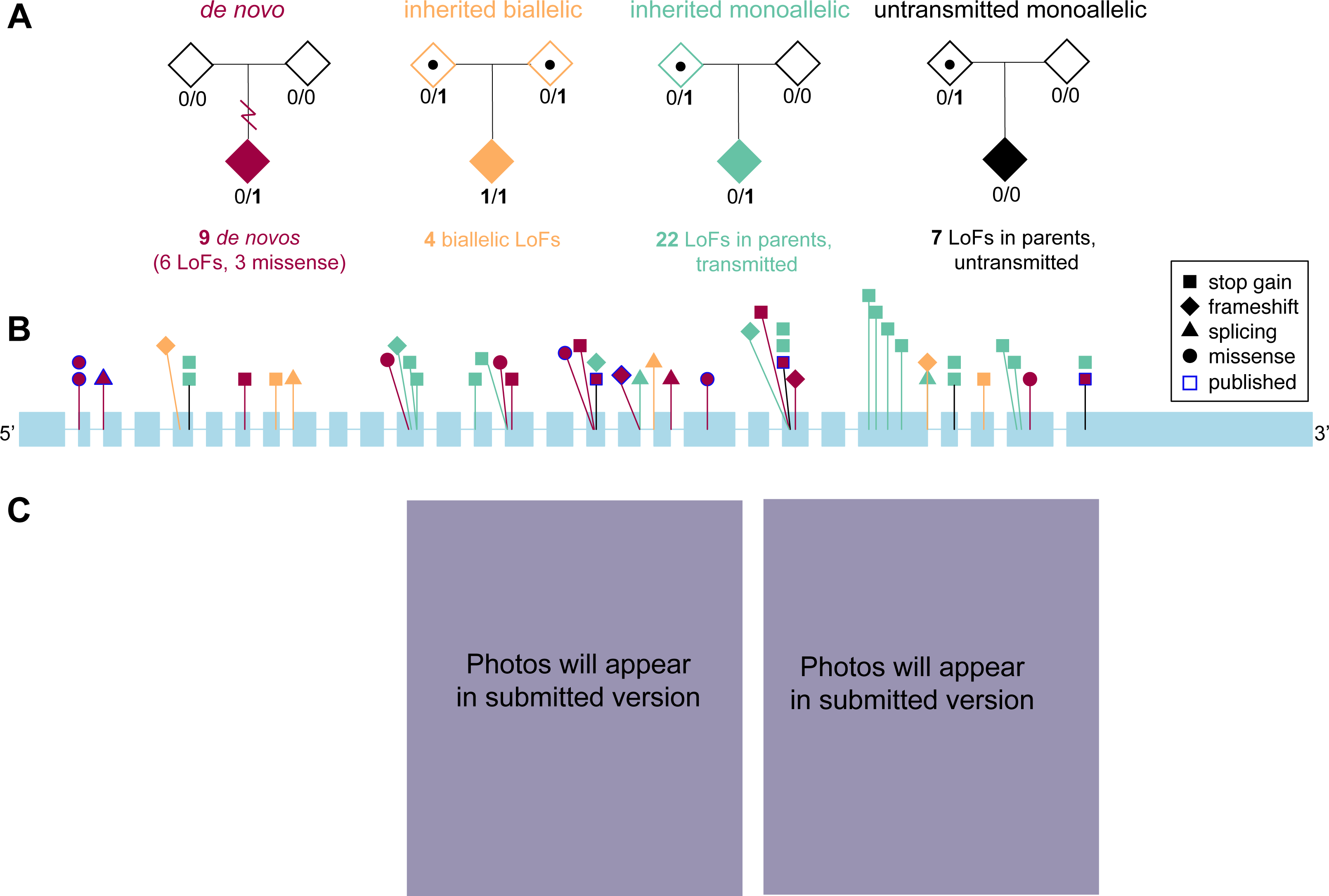
a) Summary of the damaging variants we found in *KDM5B* by mode of inheritance. b) Positions of likely damaging variants found in this and previous studies in the longest annotated transcript of *KDM5B*, ENST00000367264.2, with introns not to scale. Colours correspond to those shown in (a). There are no obvious differences in the spatial distribution of *de novo* versus monoallelic or biallelic inherited LoFs within the gene, so it is does not seem that some are less likely to be truly LoF. The points with blue borders indicate the *de novo* mutations that had been previously reported in other studies. Two large deletions are not shown (one in a biallelic proband, another of unknown inheritance). All variants are listed in Supplementary Table 5. c) Anterior-posterior facial photographs of two of the individuals with biallelic *KDM5B* variants demonstrating narrow palpebral fissures, dark eyelashes, smooth philtrum and a thin upper vermillion border. Other affected individuals shared these features. Informed consent was obtained to publish these photographs.

The four individuals with biallelic *KDM5B* variants have ID and variable congenital abnormalities (Supplementary Table 4), in line with those seen in other disorders of the histone machinery ^29^. Affected individuals have a distinctive facial appearance with narrow palpebral fissures, arched or thick eyebrows, dark eyelashes, a low hanging columella, smooth philtrum and a thin upper vermillion border (e.g. Figure 4C). Structural abnormalities observed were agenesis of the corpus callosum and a cardiac defect each in one individual. However, in contrast to other disorders of the histone machinery where growth is often promoted or restricted, there was no consistent growth pattern. Other than ID, there were no consistent phenotypes or distinctive features shared between the biallelic and monoallelic individuals, or within the latter group. Of the 26 probands with inherited LoFs in *KDM5B*, five of them were reported to have a parent who had at least one clinical phenotype shared with the child (two mothers, three fathers). However, for only two families was this the parent who carried the LoF. There was no evidence for a parent-of-origin bias in which parent transmitted the LoF. Thus, the reason for the apparent incomplete penetrance of *KDM5B* LoF variants warrants further investigation.

## Discussion

We found the contributions of *de novo* and recessive coding variants were approximately equal among DDD probands with Pakistani ancestry (both ~30%), but recessive causes contribute less than a tenth of the disease burden of dominant causes in the probands with European ancestry (3.6% vs 49.9% respectively). We showed that this difference between ancestries is entirely accounted for by differing levels of autozygosity resulting from parental consanguinity in these two populations, rather than ancestry per se. Differences in autozygosity also accounted for the differing relative contributions of homozygous and compound heterozygous genotypes to recessive causation in the two populations. We estimated that about half the recessive burden could be accounted for by genes that are already known to be DD-associated. While it has been hypothesised that there are many more recessive DD-associated genes to be discovered ^6,30^, our analyses strongly suggest that the cumulative impact of these future discoveries on diagnostic yield will be modest in outbred populations, but much more substantial in populations with higher levels of parental consanguinity.

Our use of DDD parental allele frequencies allowed us to carry out this properly calibrated burden analysis for the first time (Supplementary Figure 2), but does have caveats. Any enrichment in damaging coding variants in DDD parents compared to the general population will cause us to overestimate the population frequency of such variants, as will the systematic difference between the true allele frequency of a very rare variant and its estimate from a finite sample size ^18^. Together, these effects could cause us to slightly underestimate the overall burden. Reassuringly, our estimate in the PABI subset (30.9%) is close to the 31.5% reported by genetics clinics in Kuwait ^31^, where there is a similar level of consanguinity ^32^; this is one of the few comparable quantitative estimates of the contribution of recessive coding causes to rare disease. Our results on the EABI probands are consistent with the fact that consanguinity is very rare amongst people with European ancestry. The proportion of all DD patients in the British Isles with a recessive coding cause is probably also higher than our estimate because some recessive DDs are more easily diagnosed through current diagnostic practice than dominant ones, and therefore are less likely to be recruited to a research study. For example, a particular family history may prompt clinicians to consider recessive disorders, and recessive disorders of metabolism can often be diagnosed via biochemical testing. Nevertheless, it is striking that even in the groups thought likely to have a recessive cause (probands with affected siblings or high autozygosity), we find that recessive coding variants explain fewer than 50% of patients (Figure 2).

Our results have important clinical potential because they provide data that can improve recurrence risk counselling. In the absence of a molecular diagnosis, clinical geneticists typically give empiric recurrence risk estimates based on studies with small sample sizes (e.g. ^33–35^). There is a pressing need for large-scale studies to improve these empiric risk estimates. The results from these studies could then be combined with our findings about genetic architecture to enable clinicians to give families without an exome diagnosis more tailored estimates of recurrence risk.

Recent papers have described the population structure and characteristics of South Asian ^36^ and Middle Eastern ^37^ populations and highlighted their potential for recessive disease gene discovery. Despite this expectation, and the substantially higher burden of recessive causation in the PABI subset versus EABI (Figure 2), they contributed little to our new gene discovery. This was partially due to modest sample size but was also exacerbated by the consistent overestimation of rare variant frequencies described above. Given the strong population structure in South Asia ^38^, it will be essential to have large, appropriately ancestry-matched control sets in future studies. Studies in highly consanguineous populations would also allow investigation of the different ways that autozygosity may contribute to risk of rare genetic disorders. We previously showed that high autozygosity was significantly associated with lower risk of having a pathogenic *de novo* coding mutation in a known gene for DD ^5^. This association was still significant (p=0.003) once we controlled for the presence of at least one likely damaging biallelic genotype and other known factors (Methods). This suggests that autozygosity may increase the risk of DD via mechanisms other than a single homozygous coding variant, such as through the cumulative effect of multiple coding and/or noncoding variants. However, since overall autozygosity and the number of biallelic coding variants are correlated, it is difficult to disentangle these.

Neither of the new genome-wide significant genes we discovered in this analysis (*EIF3F* and *KDM5B*) would have been found by the traditional approach of collecting unrelated patients with the same highly recognisable disorder, because damaging biallelic genotypes in these genes result in nonspecific and heterogeneous phenotypes. It is possible they could have been identified in large consanguineous families, although the *EIF3F* variant is much rarer in South Asians than non-Finnish Europeans in ExAC, so would be harder to find in the former population. In addition to its heterogeneous presentation, *KDM5B* is also unusual for a recessive gene because heterozygous LoFs appear to be be pathogenic with incomplete penetrance. Several *de novo* missense and LoF mutations in *KDM5B* had previously been reported in individuals with autism or ID ^39–41^, but LoFs had also been observed in unaffected individuals ^40^. Disorders of the histone machinery normally follow autosomal dominant inheritance with *de novo* mutations playing a major role ^29^, so it is surprising that so many apparently unaffected DDD parents carry LoFs in *KDM5B* (Figure 4). The other genes encoding H3K4 methylases and demethylases reported to cause dominant DD ^28^ all have a pLI score >0.99 and a very low pRec, in stark contrast to *KDM5B* (pLI=5×10^−5^; pRec>0.999). LoFs in some other dominant ID genes appear to be incompletely penetrant ^42^, as do several microdeletions ^43^. So far, the evidence suggests that biallelic LoFs in *KDM5B* are fully penetrant in humans, but interestingly, the homozygous knockout does show incomplete penetrance in mice, with only one strain presenting with neurological defects ^44,45^.

There are other examples of DD genes that show both biallelic and monoallellic inheritance, such as *NALCN ^47–49^*, *MAB21L2 ^50^, ITPR1 ^51,52^*, *ROR2 ^46^* and *NRXN1 ^53,54^*. In *NALCN*, *MAB21L2* and *ITPR1*, heterozygous missense variants are thought to be activating or dominant-negative. In *ROR2* and several other genes, heterozygous LoFs in downstream exons escape nonsense-mediated decay and are dominant, whereas LoFs and missense mutations in upstream exons are recessive and result in a different phenotype ^46^. The situation in *KDM5B* is distinct from these other genes: we see biallelic LoFs and *de novo* LoFs distributed throughout the gene (Figure 4), and *de novo* missense mutations that do not obviously cluster in the protein (p=0.437; method described in ^1^). *NRXN1* is more similar: biallelic LoFs cause Pitt-Hopkins-like syndrome type 2 ^54^, which involves severe ID, whereas heterozygous deletions have been shown to predispose to a broad spectrum of neuropsychiatric disorders ^53–59^ with reduced penetrance and mild or no ID, but also to cause severe ID ^57^. Until further studies clarify the true inheritance pattern of *KDM5B*-related disorders, caution should be exercised when counselling families about the clinical significance of heterozygous variants in this gene.

In summary, we have shown for the first time that recessive coding variants make only a minor contribution to severe undiagnosed DD in EABI patients, but a much larger contribution in PABI patients. We have also identified two new genome-wide significant genes for DD (*EIF3F* and *KDM5B*). Our results suggest that identifying all the recessive DD genes would allow us to diagnose a total of 5.2% of the EABI+PABI subset of DDD, whereas identifying all the dominant DD genes would yield diagnoses for 48.6%. The high proportion of unexplained patients even amongst those with affected siblings or high consanguinity suggests that future studies should investigate a wide range of modes of inheritance including oligogenic and polygenic inheritance as well as noncoding recessive variants.

## Online Methods

### Family recruitment

Family recruitment has been described previously ^4^. 7,832 trios from 7,448 families and 1,791 patients without parental samples were recruited at 24 clinical genetics centres within the United Kingdom National Health Service and the Republic of Ireland. Families gave informed consent to participate, and the study was approved by the UK Research Ethics Committee (10/H0305/83, granted by the Cambridge South Research Ethics Committee and GEN/284/12, granted by the Republic of Ireland Research Ethics Committee). The patients were systematically phenotyped: detailed developmental phenotypes were recorded using Human Phenotype Ontology (HPO) terms ^60^, and growth measurements, family history, developmental milestones etc. were collected using a standard restricted-term questionnaire within DECIPHER ^61^. DNA was collected from saliva samples obtained from the probands and their parents, and from blood obtained from the probands, then samples were processed as previously described^1^.

### Exome sequencing and variant quality control

Exome sequencing, alignment and calling of single-nucleotide variants and small insertions and deletions was carried out as previously described ^5^, as was the filtering of *de novo* mutations. For the analysis of biallelic genotypes, we chose thresholds for genotype and site filters to balance sensitivity (number of retained variants) and specificity (as assessed by Mendelian error rate and transition/transversion ratio). We removed sites with a strand bias test p-value <0.001. We then set individual genotypes to missing if they had genotype quality < 20, depth < 7 or, for heterozygous calls, a p-value from a binomial test for allele balance < 0.001. Since the samples had undergone DNA capture with either the Agilent SureSelect Human All Exon V3 or V5 kit, we subsequently only retained sites that passed a missingness cutoff in both the V3 and the V5 samples. We found that, after setting a depth filter, the proportion of missing genotypes allowed had a more substantial effect on the number of Mendelian errors than genotype quality and allele balance cutoffs (Supplementary Figure 10). Thus, we ran the biallelic burden analysis on two different callsets, using a 10% (strict) or a 50% (lenient) missingness filter, and found that the results were very similar. We report results from the more lenient filter in this paper, since it allowed us to include more variants. Genotypes were set to missing for a trio if there was a Mendelian error, and variants were removed if more than one trio had a Mendelian error and if the ratio of trios with Mendelian errors to trios carrying the variant without a Mendelian error was greater than 0.1. If any of the individuals in a trio had a missing genotype at a variant, all three individuals were set to missing for that variant.

Variants were annotated with Ensembl Variant Effect Predictor ^62^ based on Ensembl gene build 83, using the LOFTEE plugin. The transcript with the most severe consequence was selected. We analyzed three categories of variant based on the predicted consequence: (1) synonymous variants; (2) loss-of-function variants (LoFs) classed as “high confidence” by LOFTEE (including the annotations splice donor, splice acceptor, stop gained, frameshift, initiator codon and conserved exon terminus variant); (3) damaging missense variants (i.e. those not classed as “benign” by PolyPhen or SIFT, with CADD>25). Variants were also annotated with MAF data from four different populations of the 1000 Genomes Project ^63^ (American, Asian, African and European), two populations from the NHLBI GO Exome Sequencing Project (European Americans and African Americans) and six populations from the Exome Aggregation Consortium (ExAC) (African, East Asian, non-Finnish European, Finnish, South Asian, Latino), and an internal allele frequency generated using unaffected parents from the DDD.

### Ancestry inference

We ran a principal components analysis in EIGENSOFT ^64^ on 5,853 common exonic SNPs defined by the ExAC project. We set genotypes with GL<20 to missing and excluded SNPs with >2% missingness, and then excluded samples with >5% missingness from this and all subsequent analyses. We calculated principal components in the 1000 Genomes Phase III samples and then projected the DDD samples onto them. We grouped samples into three broad ancestry groups (European, South Asian, and Other) as shown in Supplementary Figure 1 (right hand plots). By drawing ellipses around the densest clusters of DDD samples, we defined two narrower groups: European Ancestry from the British Isles (EABI) and Pakistani Ancestry from the British Isles (PABI).

For the burden and gene-based analysis, we primarily focused on these narrowly-defined EABI and PABI groups because it is difficult to accurately estimate population allele frequencies in more broadly defined groups. For example, in 4,942 European-ancestry probands, the number of observed biallelic synonymous variants was slightly higher than the number of expected (ratio = 1.06; p=2.7×10^−4^).

### Calling autozygous regions

To call autozygous regions, we ran bcftools/roh^65^ (bcftools version 1.5-4-gb0d640e) separately on the different broad ancestry groups. We LD pruned our data to avoid overcalling small runs of homozygosity as autozygous regions. Because rates of consanguinity differ dramatically between EABI and PABI, we chose r^2^ cutoffs for each that brought the ratio of observed to expected biallelic synonymous variants with MAF<0.01 closest to 1 (see below for calculation of the number expected): PLINK options ‐‐indep-pairwise 50 5 0.4 for EABI and ‐‐indep-pairwise 50 5 0.8 for PABI.

### Defining sample subsets

We stratified probands by high autozygosity (>2% of the genome classed as autozygous), whether or not they had an affected sibling, and whether or not they already had a likely diagnostic dominant or X-linked exonic mutation (a likely damaging *de novo* mutation or inherited damaging variant in a known monoallelic DDG2P gene (<u>http://www.ebi.ac.uk/gene2phenotype/</u>) [if the parent was affected] or a damaging X-linked variant in a known X-linked DDG2P gene). The 4,458 patients who had no such diagnostic variants were included in the “undiagnosed” set, along with 193 patients who had biallelic genotypes in recessive DDG2P genes or potentially diagnostic variants in monoallelic or X-linked

DDG2P genes but had high autozygosity or affected siblings. There were 1,366 EABI and 23 PABI probands in the diagnosed set, and 4,318 EABI and 333 PABI probands in the undiagnosed set. For the set of probands with affected siblings shown in Figures 1 and 2, we restricted to families from which more than one independent (i.e. non-MZ twin) child was included in DDD and in which the siblings’ phenotypes were more similar than expected by chance given the distribution of HPO terms in the full cohort (HPO similarity p-value < 0.05 ^10^).

For the burden analysis and gene-based tests, we removed 11 probands with uniparental disomy, and one individual from every pair of probands who were related (kinship > 0.044, estimated by PCRelate ^66^, equivalent to third-degree relatives). We also removed 924 parents reported to be affected, since one might expect these to be enriched for damaging variants compared to the general population, and 9 European parents with an abnormally high number of rare (MAF<1%) synonymous genotypes (>834, compared to the 99.9^th^ percentile of 223), but we retained their offspring.

### Burden analyses and gene-based tests

Variants were filtered on class (LoF, damaging missense or synonymous) and by different MAF cutoffs. Variants failing the MAF cutoff in any of the publicly available control populations, the full set of unaffected DDD parents, or the unaffected DDD parents in that population subset (PABI or EABI) were removed.

Following the approach we used previously ^10^, we calculated *B_g,c_,* the expected number of rare biallelic genotypes of class *c* (LoF, damaging missense or synonymous) in each gene *g*, as follows:

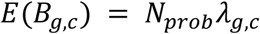

where *N_prob_* is the number of probands *λ_g,c_* and is the expected frequency of biallelic genotypes of class *c* in gene, calculated as follows:

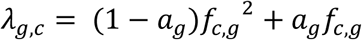

where *f_c,g_* is the cumulative frequency of variants of class *c* in gene *g* with MAF less than the cutoff, and *a_g_* is the fraction of individuals autozygous at gene *g*. An individual was defined as being autozygous if he/she had a region of homozygosity with any overlap of gene *g*; in practice, autozygous regions almost always overlapped genes completely rather than partially.

The rate of LoF/damaging missense compound heterozygous genotypes is:

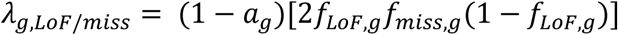

To calculate the cumulative frequency of variants of class *c* in gene *g*, *f_c,g_*, we first phased the variants in the parents based on the inheritance information. The cumulative frequency is then given by:

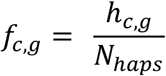

where *h_c_* is the number of parental haplotypes with at least one variant of class *c* in gene *g*, and *N_haps_* is the total number of parental haplotypes.

For each gene, we calculated the binomial probability (given *N_prob_* probands and rate*λ_g,c_*) of the observed number of biallelic genotypes of class *c.* We did this for four consequence classes (biallelic LoF, biallelic LoF+LoF/damaging missense, biallelic damaging missense, and biallelic LoF+LoF/damaging missense+biallelic damaging missense) and for two sets of probands (EABI only, and EABI+PABI). We did not analyze PABI separately due to low power.

For the set of EABI only, we conducted a simple binomial test. For the combined EABI+PABI test, we took into account the different ways in which *n* or more probands with the relevant genotype could be distributed between the two groups and the probability of observing each combination using population-specific rates (e.g. two observed biallelic genotypes could be both seen in EABI, both in PABI, or one in each). We then summed these probabilities across all possible combinations to obtain an aggregate probability for sampling *n* or more probands by chance, as described in ^10^.

For some genes, *λ_g,c_* was estimated to be 0 in one or both populations because there were no variants in the parents that passed filtering. The vast majority of these also had *O*(*B_g,c_*) = 0. We dropped these genes from the tests, but still included them in our Bonferroni correction. We also excluded 715 genes either because they were in the HLA region or because they were classed as having suspiciously many or suspiciously few synonymous or synonymous+missense variants in ExAC, leaving 18,630 genes. We thus set a significance threshold of 0.05/(8 tests × 18,630 genes) = p<3.4×10^−7^. For Supplementary Figure 6, we ordered the genes by their lowest p-value, randomized the order of genes with the same p-value, then tested for a difference in the distribution of ranks between recessive DDG2P genes and all other genes using a Kolmogorov-Smirnov test.

For the burden analysis, we summed up the observed and expected number of biallelic genotypes across all genes to give *O(B_c_)* and *E(B_c_)*, then calculated their difference *O(B_c_)*–*E(B_c_)* (the excess) and their ratio 
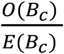
. Under the null hypothesis, we expect *O(B_c_)* to follow a Poisson distribution with rate *E(B_c_)*.

### Alternative methods for estimating the expected number of biallelic genotypes

For Supplementary Figure 2, we used two alternative methods to the one described above for estimating the expected number of biallelic genotypes. Firstly, we calculated the cumulative frequency based on ExAC by summing the frequencies of individual variants of class *c* in gene *g* separately for non-Finnish Europeans (NFE) for EABI and from South Asians (SAS) for PABI:

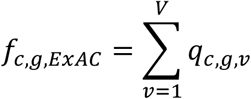

For this, we restricted to ExAC variants within the intersection of the V3 and V5 Agilent probes used in DDD (including 100bp flanks on each side), and removed variants if they had >50% missingness in the relevant ExAC population.

We also used a modified version of the method described by Jin *et al.* ^3^ for approximating the expected number of biallelic genotypes by fitting a polynomial regression on the mutability:

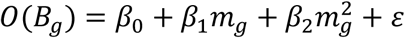

where *O(B_g_)* is the observed number of biallelic genotypes for gene *g,m_g_* is the mutability for gene *g* calculated using the method of Samocha *et al.* ^67^, *β_0_,β_1_* and *β_2_* are coefficients, and *ε*is a random noise term. We fitted this model to the observed number of biallelic synonymous genotypes (MAF<0.01) for each gene to estimate *β_0_,β_1_* and *β_2_*, then calculated the expected number of biallelic genotypes for each gene using the fitted value:

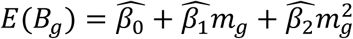

Note that we did not do the additional step described in Jin *et al.* in which they calculate the expected number of biallelic genotypes for gene *g* as 
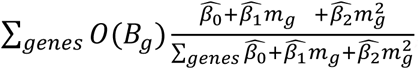
, since this constrains the total expected number of biallelic genotypes to be equal to the number observed, making exome-wide burden analysis impossible.

### Estimating the proportion of cases with diagnostic biallelic coding variants or *de novo* mutations

We are interested in estimating the proportion of probands with diagnostic variants of consequence class *π_c_*. Under the null hypothesis in which none of the genotypes of class *c* are pathogenic, the number of such genotypes we expect to see in *N_pr_* probands is:

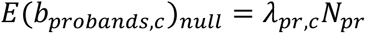

where 
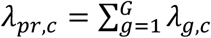
 is the total expected frequency of genotypes of class *c* across all genes.

However, under the alternative hypothesis, suppose that some fraction *φ_causal,c_* of genotypes in class *c* cause DD, and some fraction *φ_lethal,c_* are lethal. Assuming complete penetrance, we can thus split *E(b_probands,c_)* into genotypes that are due to chance and those that are diagnostic:

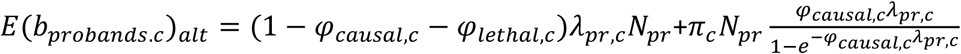

The component due to chance is 
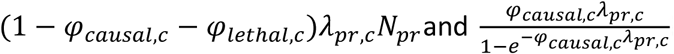
 is the average number of pathogenic genotypes per individual, given that the individual has at least one such genotype.

In *N_pa_* healthy parents, biallelic genotypes of class *c* that is not pathogenic:

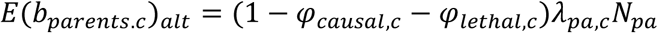

where *λ_pa,c_* is the expected rate of biallelic genotypes of class *c* in the parents, given the cumulative frequencies estimated in the same set of people, and the autozygosity rates. We can thus obtain a maximum likelihood estimate for*φ_c_* = *φ_causal,c_*+*φ_lethal,c_* using *O_pa,c_*, the observed number of biallelic genotypes of class *c* in the parents:

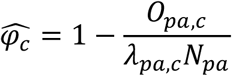

The 95% confidence interval for 
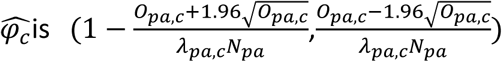
. We show estimates of *φ_c_* from the EABI and PABI parents in Supplementary Figure 5. To estimate *π_c_*, we the combined data from both populations for MAF<0.01 variants to estimate *φ_c_*, and obtained the following maximum likelihood estimates and 95% confidence intervals: *φ_LoF/LoF_* = (0.141 (0.046,0.238),*φ_LoF/miss_* = 0.083 (−0.009,0.175),and *φ_miss/miss_* = 0.007(−0.028,0.042).

For biallelic genotypes, we can substitute *φ_c_* into the expression above, substitute *O_pr,c_* for *E(b_probands_)* and rearrange to obtain a maximum likelihood estimate of *π_c_*:

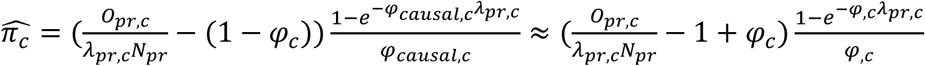

We cannot disentangle *φ_causal,c_* and *φ_lethal,c_* with the available data, but we find that the ratio 
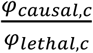
 makes very little difference to the estimate of 
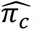
, so we make the assumption that *φ_causal,c_*= *φ_c_*. The 95% confidence interval is then 
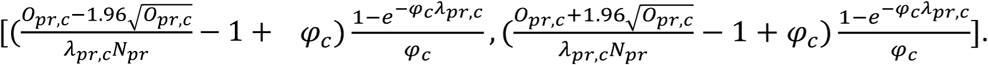
 .

*De novo* mutations were called as previously described ^5^, selecting the threshold on ppDNM (posterior probability of a *de novo* mutation) such that the observed number of synonymous *de novos* matched the number expected. Using Sanger validation data from an earlier dataset ^1^, we adjusted the observed number of mutations to account for specificity and sensitivity as follows:

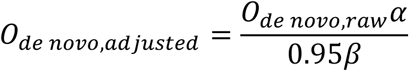

where *α* is the positive predicted value (the proportion of candidate mutations that are true positive at the chosen threshold) and *β* is the sensitivity to true positives at the same threshold. The adjustment by 0.95 is due to exome sequencing being only about 95% sensitive. The overall *de novo* mutation rate *λ_de novo,c,pr_* was calculated in different sets of probands using the model from ^67^, adjusting for sex, as described previously ^5^ .

Since we cannot estimate *φ_c_* for de novo mutations using the parents, as we did for recessive variants, we instead set *φ_DN LoF_* to 0.099, the fraction of genes with pLI>0.99. This estimate is more speculative than the directly observed depletion of biallelic genotypes above, but we note that the estimate of *π_DN LoF_* for the full set of 7,832 trios only increases from ~0.129 to ~0.154 if we increase *φ_DN LoF_* from 0.01 to 0.3. To estimate *φ_DN missense_*, we make use of this relationship:

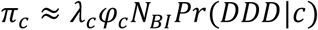

where *N_BI_* is the population of the British Isles, and *Pr*(*DDD|c*) is the probability that an individual is recruited to the DDD given he/she has a pathogenic mutation of class *c.* If we assume that this recruitment probability is the same for *de novo* missense mutations as for *de novo* LoFs, we can write:

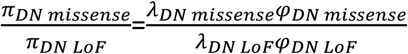

We know *λ_DN missense_* and *λ_DN LoF_*, will assume *φ_DN LoF_* = 0.099 so we can estimate *π_DN LoF_*, and can thus write the number of *de novo* missense mutations we expect to see as:

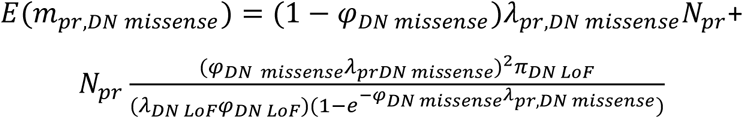

We calculated *E*(*m_pr,DN missense_*) for a range of values of *φ_DN missense_* and found that *φ_DN missense_* = 0.036 best matched the observed data, so we used this value for estimating *π_DN missense_*.

### Effect of autozygosity on risk of having a diagnostic *de novo*

We fitted a logistic regression on all EABI and PABI probands as follows:

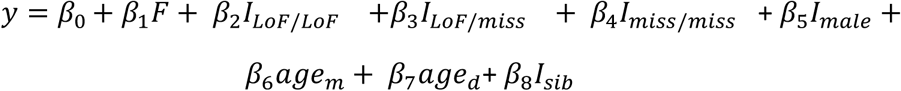

where *y*is an indicator for having a diagnostic *de novo* coding mutation (1) or not (0), *F* is the overall fraction of the genome in autozygous segments, *I_LoF/LoF_*, *I_LoF/miss_* and *I_miss/miss_* are indicators for the presence of at least one biallelic genotype in the relevant class, *age_m_* and *age_d_* are the parental ages at birth for the mother and father respectively,and *I_male_* and *I_sib_* are indicators for being male and having an affected sibling respectively. In this joint model, the significant covariates were *F*(*β_1_* =−10.96; p=0.003), *I_LoF/LoF_*(*β_2_*= −0.92; p=3x10^−4^), *I_male_*(*β_5_*=−0.38; p=5x10^−10^), *age_d_*(*β_7_*=0.017; p=0.005) and *I_sib_*(*β_8_*=−0.87; p=1x10^−11^). The autozygosity effect is equivalent to a ~2-fold decreased chance of having a diagnostic *de novo* for a DDD patient who is offspring of first cousins (expected autozygosity= 6.25%) .

### Structural analysis of EIF3F

Human EIF3:f (pdb 3j8c:f) was submitted to the Protein structure comparison service PDBeFold at the European Bioinformatics Institute ^68,69^. Of the close structural matches returned, the X-ray yeast structure pdb entry 4OCN was chosen to display the human variant position, as the structural resolution (2.25Å) was better than the human EIF3:f pdb 3j8c:f structure (11.6Å) and it was the most complete structure among the yeast models. In order to map the Phe232 variant onto the equivalent position on the yeast structure, the structural alignment from PDBeFold was used. Solvent accessibility was calculated using the Naccess software ^70^ using the standard parameters of a 1.4Å probe radius. Amino acid sequence conservation was calculated using the Scorecons server ^71^ and displayed using sequence logos ^72^.

### Validation of *KDM5B* variants by targeted re-sequencing

We re-sequenced all *KDM5B de novo* mutations and inherited LoF variants, with the exception of two large deletions. PCR primers were designed using Primer3 to amplify the site of interest, generating approximately a 230 bp product centred on the site. PCR amplification of the targeted regions was carried out using JumpStart^TM^ AccuTaq^TM^ LA DNA Polymerase (Sigma-Aldrich), using 40ng of input DNA from the proband and their parents. Unique identifying tag sequences were introduced into the PCR amplicons in a second round of PCR using KAPA HiFi HotStart ReadyMixPCR Kit (KapaBiosystems). PCR amplicons were pooled and 96 products were sequenced in one MiSeq lane using 250bp paired-end reads. Reference and alternate read counts extracted from the resulting bam files and were used determine the presence of the variant in question. In addition, read data were visualised using IGV.

### Transmission-disequilibrium test on *KDM5B* LoFs

We observed 15 trios in which one parent transmitted a LoF to the child, 5 trios in which one parent had a LoF that was not transmitted, 2 quartets in which one parent had a LoF that was transmitted to one out of two affected children, and 4 trios in which both parents transmitted a LoF to the child. We tested for significant over-transmission using the transmission-disequilibrium test as described by Knapp ^73^. There were 7 LoFs (including one large deletion) observed in probands whose parents were not originally sequenced, which we excluded from the TDT. Of the six for which we attempted validation and segregation analysis, one was found to be *de novo* and five inherited.

### Searching for coding, regulatory or epigenetic modifiers of *KDM5B*

We defined a set of genes that might modify *KDM5B* function as: interactors of *KDM5B* obtained from the STRING database of protein-protein interactions ^74^ (*HIST2H3A*, *MYC*, *TFAP2C*, *CDKN1A*, *TFAP2A*, *SETD1A*, *SETD1B*, *KDM1A*, *KDM2B*, *PAX9*) plus those mentioned by Klein *et al.* ^75^ (*RBBP4*, *HDAC1*, *HDAC4*, *MTA2*, *CHD4*, *FOXG1*, *FOXC9*), as well as all lysine demethylases, lysine methyltransferases, histone deacetylases, and SET domain-containing genes from <u>http://www.genecards.org/</u>. The final list contained 95 genes. We looked for LoF or rare missense variants in these genes in the monoallelic *KDM5B* LoF carriers that might have a modifying effect, but found none that were shared by more than two of the *de novo* carriers.

We also looked for indirect evidence of a regulatory “second hit” near *KDM5B* by examining the haplotypes of common SNPs in the region (Supplementary Figure 8). DDD probands and a subset of their parents were genotyped on either the Illumina OmniExpress chip or the Illumina CoreExome chip. We performed variant and sample quality control for each dataset separately. Briefly, we removed variants and samples with high data missingness (>=0.03), samples with high or low heterozygosity, sample duplicates, individuals of African and East Asian ancestry, and SNPs with MAF<0.005. We then ran SHAPEIT2 ^76^ to phase the SNPs within 2Mb either side of *KDM5B*. To make Supplementary Figure 8, we used the 

~~~
heatmap()
~~~

 function in R to cluster the phased haplotypes using the default hierarchical clustering method (based on Euclidean distance).

We looked at methylation levels in the *KDM5B* LoF carriers to search for an “epimutation” (hypermethylation on or around the promoter) that might be acting as second hit. DNA from 64 DDD whole blood samples comprising 41 probands with a *KDM5B* variant and 23 negative controls was run on an Illumina EPIC 850K methylation array. Negative controls were selected from DDD probands with *de novo* mutations in genes not expressed in whole blood (*SCN2A*, *KCNQ2*, *SLC6A1*, and *FOXG1*), since we would not expect these to significantly impact the methylation phenotype in that tissue. Samples were randomised on the array to reduce batch effects, and were QCed using a combination of data from control probes and numbers of CpGs that failed to meet the standard detection p-value of 0.05. Based on these criteria, two samples failed and were excluded from further analysis (one of the negative controls and one of the inherited *KDM5B* LoF carriers). We analyzed a subset of CpGs in and around the *KDM5B* promoter region: the CpG island in the *KDM5B* promoter itself, and a CpG island in the promoter of *KDM5B-AS1*, a lnc-RNA not specifically associated with *KDM5B*, but also highly expressed in the testis. We also extended analysis 5kb on either side of the start and stop sites of the *KDM5B* promoter. We examined the distribution of the beta values (the ratio of methylated to unmethylated alleles) at each of the CpGs in the 10kb region (Supplementary Figure 8).

## Acknowledgements

We thank the families for their participation and patience. We are grateful to the Sanger Human Genome Informatics team, the Sample Management team, the Illumina High-Throughput team, the New Pipeline Group team, the DNA pipelines team and the Core Sequencing team for their support in generating and processing the data. We also thank Petr Danacek for help with calling the regions of homozygosity, and Kaitlin Samocha for useful discussions and comments on the manuscript and for providing the mutability estimates for Ensembl transcripts. The DDD study presents independent research commissioned by the Health Innovation Challenge Fund (grant HICF-1009-003), a parallel funding partnership between the Wellcome Trust and the UK Department of Health, and the Wellcome Trust Sanger Institute (grant WT098051). The views expressed in this publication are those of the author(s) and not necessarily those of the Wellcome Trust or the UK Department of Health. The study has UK Research Ethics Committee approval (10/H0305/83, granted by the Cambridge South Research Ethics Committee and GEN/284/12, granted by the Republic of Ireland Research Ethics Committee). The research team acknowledges the support of the National Institutes for Health Research, through the Comprehensive Clinical Research Network.

## Author Contributions

Exome sequence data analysis: H.C.M., J.F.M. Protein structure modelling: J.S., Methylation analysis: J.H., E.R. Clinical interpretation: W.D.J. Data processing: G.G., M.N., J.K., C.F.W. Experimental validation: E.P. Methods development: H.C.M., P.S., M.E.H., J.C.B. Data interpretation: H.C.M., J.S., J.H., N.A., M.E.H., J.C.B. Patient recruitment and phenotyping: M.B., J.D., R.H., A.H., D.S.J., K.J., D.K, S.A.L., S.G.M., J.M., M.J.P., M.S., P.D.T., P.C.V., and M.W. Experimental and analytical supervision: C.F.W., D.R.F., H.V.F., M.E.H., J.C.B. Project Supervision: J.C.B. Writing: H.C.M., W.D.J., J.S., J.H., J.C.B.

## Supplementary Material

**Supplementary Figure 1**: Principal components analysis of the 1000 Genomes Phase 3 samples (left) with DDD samples projected on top of them (right). The ellipses used to define the EABI and PABI populations in DDD are shown on the PC2 versus PC3 plot.

**Supplementary Figure 2**: Number of observed and expected biallelic synonymous genotypes per individual, estimated across 5684 EABI and 356 PABI probands. The grey lines indicate the observed number and the points with small black lines represent the expected number with a 95% confidence interval. Two-sided p-values are indicated. The expected numbers were calculated in different ways as described in the Methods. Note that the estimate based on the DDD parental haplotypes (black points) best matches the observed data, and that use of the ExAC frequencies or the polynomial model of Jin *et al. ^3^* gives estimates of the expected number that are significantly different from the observed number.

**Supplementary Figure 3**: Histograms of levels of autozygosity across EABI and PABI probands.

**Supplementary Figure 4**: A) Burden (ratio of observed to expected) or B) excess (observed-expected) of biallelic genotypes in EABI undiagnosed probands for different MAF cutoffs. The coloured lines show 95% confidence intervals. The points for different consequence classes at the same MAF cutoff have been slightly scattered along the x-axis for ease of visualisation. Note that we do not show results for the PABI subset, because of inaccurate allele frequency estimates in this small sample.

**Supplementary Figure 5**: Estimates of φ, the proportion of biallelic genotypes that are lethal or cause DD. These were estimated from the parental data. See Methods for details. The points show maximum likelihood estimates and the lines show 95% confidence intervals. Points at the same MAF cutoff have been slightly scattered along the x-axis for ease of visualisation.

**Supplementary Figure 6**: Distribution of the ranks of minimum p-values per gene for known recessive genes versus all other genes. The order of genes with the same minimum p-value was randomised. A Kolmogorov-Smirnov (KS) test indicated that these distributions were significantly different.

**Supplementary Figure 7**: Anterior-posterior facial photographs of individuals with the homozygous Phe232Val variant in EIF3F. DECIPHER IDs are shown in the top right corner. Affected individuals did not have a distinctive facial appearance. Individual 265452 (leftmost) had muscle atrophy, as demonstrated in photographs of the anterior surface of the hands which show wasting of the thenar and hypothenar eminences.

**Supplementary Figure 8**: Plot showing haplotypes of common SNPs around *KDM5B* in individuals with *de novo* missense or LoF mutations or with monoallelic or biallelic LoFs. These is no evidence for a local haplotype shared by multiple probands with monoallelic LoFs that was not also present in an unaffected parent with a monoallelic LoF. The region shown lies between two recombination hotspots. The rows represent phased haplotypes, with orange and green rectangles corresponding to the different alleles at the SNPs at the positions indicated along the bottom. Hierarchical clustering has been applied to the haplotypes, as indicated by the dendrogram on the left, and the labels on the right indicate which individual carries the haplotype, and whether the individual was a proband carrying a *de novo* (purple), a biallelic LoF (dark green), or an inherited heterozygous LoF (yellow), or a parent carrying a heterozygous LoF (pink).

**Supplementary Figure 9**: Violin plots of the beta values (the ratio of methylated to unmethylated alleles) at each of the CpGs in the 10kb region around the *KDM5B* promoter. The CpGs within the *KDM5B* and *KDM5B-AS1* promoters are annotated below the plot, with coordinates relative to hg19. The bottom panel shows the negative controls (probands with likely causal *de novo* mutations in known DD genes not expressed in blood), and the other panels show probands with variants in *KDM5B* that are either biallelic (top panel), *de novo* (second panel) or monoallelic and inherited (third panel).

**Supplementary Figure 10**: Plots showing effect of variant filtering strategies on number of variants, Mendelian errors and Ti/Tv. We first set genotypes to missing based on genotype quality (GQ), depth (AD) and the p-value from a test of allele balance (p_AB_), and then removed sites according to the proportion of missing genotypes.

**Supplementary Table 1**: Estimates of the π, the proportion of probands explained by diagnostic biallelic coding genotypes or *de novo* coding mutations, for different sample sets. Shown are the maximum likelihood estimates for π, and a 95% confidence interval. See Methods for how these were calculated.

**Supplementary Table 2**: Results from tests of an excess of damaging biallelic genotypes for all genes. The lowest p-value out of the eight tests conducted for each gene is shown. We give results for the stringent ancestry filter (4318 EABI probands and 333 PABI probands), as shown in Table 1, as well as the lenient ancestry filter (4942 European ancestry probands and 498 South Asian ancestry probands).

**Supplementary Table 3**: Phenotypes of the nine patients homozygous for the *EIF3F* Phe232Val variant.

**Supplementary Table 4**: Phenotypes of the four probands with biallelic *KDM5B* variants.

**Supplementary Table 5**: *De novo* mutations in *KDM5B* from this and previous studies, and inherited LoFs in *KDM5B* from this study.

